# Heterospecific physical interference modulates the reproductive attributes of a ladybird beetle

**DOI:** 10.1101/2023.05.23.541875

**Authors:** Mohd Sariq, Omkar, Geetanjali Mishra

**Affiliations:** Ladybird Research Laboratory, Department of Zoology, University of Lucknow, Lucknow- 226007, India

**Keywords:** *Propylea dissecta*, *Cheilomenes sexmaculata*, reciprocal mating interaction, reproductive interference, reproductive fitness

## Abstract

*Propylea dissecta* and *Cheilomenes sexmaculata* (Coleoptera: Coccinellidae), are similar sized, co-ocurring aphidophagous insects that share common resources. We conducted the current study to observe the phenomenon of reproductive interference and effect of presence of heterospecific adults (*C. sexmaculata*) on the reproductive output of *P. dissecta*. For this, we used two reciprocal mating combinations of heterospecifics. The first heterospecific combination used *P. dissecta* males with *C. sexmaculata* females and the second heterospecific mating combination used *P. dissecta* females with *C. sexmaculata* males. To further examine the effect of *C. sexmaculata* male on *P. dissecta* female, we used mixed mating treatments (*C. sexmaculata* male with conspecific pair of *P. dissecta*). Our results suggested that (1) in the first heterospecific mating combination, mating did not commence between *P. dissecta* male *C. sexmaculata* female, while in the second heterospecific mating combination, mating was recorded between *P. dissecta* female and *C. sexmaculata* male, (2) *C. sexmaculata* male interferes the conspecific mating in *P. dissecta* through multiple mating attempts which resulted in reduced fitness and reproductive success of *P. dissecta*. Overall, we illustrated the negative effects of reproductive interference caused by *C. sexmaculata* on the reproductive output (fecundity and egg viability) of *P. dissecta*.

## Introduction

Reproductive interference may occur as a result of mating interactions among heterospecific individuals. (Gröning & Hochkirch, 2008; Burdfield-Steel & Shuker, 2011). Reproductive interference is seen in different taxa, such as fishes (Tsurui-Sato *et al*., 2019), amphibians (Hettyey & Pearman, 2000; Hettyey *et al*., 2014), birds (Jones & Hunter, 1998), mammals (Carranza *et al*., 2017) and insects (Noriyuki & Osawa, 2016; Shuker & Burdfield-Steel, 2017).

Such heterospecific reproductive interaction usually occurs when both or one of the sexes (usually females) lacks premating barriers and they are unable to reject forced copulation attempts by heterospecific mates (Ribeiro & Spielman, 1986). If postmating barriers are complete then the cost of heterospecific mating is high (Liou & Price, 1994). This mating results in waste of time, energy and gametes, and reduces the fitness of one or both species (Gröning & Hochkirch, 2008; Burdfield-Steel & Shuker, 2011). Physical damages are caused during reproductive interference, due to incompatible morphologies (Kyogoku & Nishida, 2013; Ronn *et al*., 2007; Kyogoku & Nishida, 2013; Kyogoku & Sota, 2015). All aspects of reproduction (*e*.*g*. mate finding, territoriality, frequency of conspecific courtship and mating, fecundity of females, fertility of eggs) and fitness of individuals are influenced by reproductive interference (Gröning & Hochkirch, 2008; Shuker & Burdfield-steel, 2017). Heterospecific mating is suggested to occur due to failed species recognition, and species having the same size and same mating signals might also be responsible for the misdirected mating (Gröning & Hochkirch, 2008; Burdfield-Steel & Shuker, 2011).

Such a heterospecific reproductive interaction has possibly evolved as a result of male promiscuity (Gröning & Hochkirch, 2008) or due to incomplete species recognition (Shuker & Burdfield-steel, 2017). Often in heterospecific matings inviable offspring are produced but, in some cases, viable hybrids are also produced by heterospecific mating (Villa *et al*., 2021). Reproductive interference can maintain shape and community structure, and evolutionary trajectories of population (Kyogoku & Wheatcroft, 2020) as it promotes species coexistence (Villa *et al*., 2021). Some authors suggest that reproductive interference is a form of ecological competition (Shuker & Burdfield-steel, 2017); however, it differs from competition as there is neither shared resource nor density dependence evidence (Gröning & Hochkirch, 2008).

Reproductive interference has been reported extensively in closely related congeneric species such as *Aedes aegypti* L. and *Aedes albopictus* Skuse (Paton & Bonsall, 2019), *Callosobruchus chinensis* L. and *Callosobruchus maculatus* Fabricius (Kyogoku & Nishida, 2012; Kishi, 2015), *Drosophila melanogaster* Meigen and *Drosophila simulans* Sturtevant (Sultanova *et al*., 2020), *Tribolium castaneum* Herbst and *Tribolium confusum* Jacquelin du Val (Gilad & Scharf, 2019). However, its incidence in different genera is less often reported, for example in *Lygaeus equestris* L., *Lygaeus creticus* Lucas, *Spilostethus pandurus* Scopoli and *Oncopeltus fasciatus* Dallas (Shuker *et al*., 2015), Eastern *Perithemis tenera* Say and *Tabanus spp* or *Ancyloxypha numitor* Fabricius (Schultz & Switzer, 2001).

Reproductive isolation occurs in phytophagous ladybird beetles due to interspecific mating between *Epilachna* species (Nakano & Sciences, 1994) and *Henosepilachna* species (Coccinellidae) (Hirai *et al*., 2006). Reproductive interference is known to cause niche partitioning in aphidophagous insects (Noriyuki, 2015). Ladybird beetles *Harmonia* sp. (Coccinellidae) show reproductive interference, females of both species (*Harmonia axyridis* and *H. yedoensis*) laid inviable eggs after interspecific mating, because premating barriers are absent in *Harmonia* species. With a high density of *Harmonia axyridis, Harmonia yedoensis* fails in conspecific mating (Noriyuki & Osawa, 2016). Niche partitioning not only occurs in *Harmonia* species but has also been reported in other predatory ladybird beetle, such as *Adalia* spp., *Coccinella* spp., *Chilocorus* spp., *Coccidula* spp., etc. Coexistence of species, frequency of population, home range and other different factors are responsible for interspecific mating or reproductive interference.

On the basis of the above studies, we can infer that when two or more species or genera coexist together they interact with each other for food, niches and mating partners. *Propylea dissecta* and *Cheilomenes sexmaculata* are known to co-occur and share common resources, thus we hypothesized that reproductive success of *P. dissecta* will be significantly affected by the presence of heterospecific adults of *C. sexmaculata*.

## Material and Methods

### Stock maintenance

Adults of *Propylea dissecta* (N = 50) and *Cheilomenes sexmaculata* (N = 50) were collected from agricultural fields in Lucknow, India (26°50 N, 80°54 E). Adults of both species were paired and placed in beakers separately under laboratory conditions (27 ± 2°C temperature; 65 ± 5% relative humidity; 14L:10D photoperiod in Biochemical Oxygen Demand Incubators; Yorco Super Deluxe, YSI-440 New Delhi, India) and were provided with *ad libitum* supply of aphids, *Aphis craccivora* (maintained on the host plant, *Vigna unguiculata* L.). These rearing conditions were maintained throughout the experiment. Beakers were checked daily for oviposition; eggs laid were collected. Post-hatching, larval instars were transferred in separate Petri dish (9.0 × 2.0 cm, each Petri dish size) with the help of fine tip camel hair brush.

### Experimental Setup

Each larva was placed individually in plastic Petri dishes (size as above) and was provided with aphid, *A. craccivora* until emergence. Post-emergence the adults of both species were given *ad libitum* supply of aphids. Ten-day-old unmated and unrelated adults of both *C. sexmaculata* and *P. dissecta* were used in experiment.

Ten-day-old adults of *P. dissecta* and *C. sexmaculata* were collected from laboratory stock. To see the role of reproductive interference on mating and reproductive parameters, three mating treatments were formed using conspecific and heterospecific adults (a) ten-day-old unmated male *P. dissecta* and ten-day-old unmated female *P. dissecta*, (b) ten-day-old unmated male *P. dissecta* and ten-day old unmated female *C. sexmaculata*, and (c) ten-day-old unmated male *C. sexmaculata* and ten-day-old unmated female *P. dissecta*. A fourth treatment that assessed the effect of presence of conspecific and heterospecific mates on mating and reproductive parameters was also conducted. In this mixed treatment, (d) ten-day-old unmated male and female of *P. dissecta* and an unmated male of *C. sexmaculata* (heterospecific adult) were placed in same Petri dish. All replicates were observed for 30 minutes for commencement of mating. Replicates in which mating did not commence within 30 minutes of cohabitation were discarded. 30 replicates were conducted per treatment.

Treatments (b) and (c) were representative of heterospecific mating or reciprocal mating interaction, while the fourth treatment indicated reproductive interference. In total, 30 replicates of heterospecific pairs (b) and (c) were observed for mating, out of which mating did not occur in any of the replicates of (b). However, in heterospecific pair of (c) treatment, mating was recorded in 13 replicates. Thus, 30 replicates of heterospecific pair (b) and 17 replicates of heterospecific pair (c) were discarded. Since we wanted to observe heterospecific mating and the effect of heterospecific male, we did not use pairing between *C. sexmaculata* males and females.

Mating incidence, mating parameters, *i*.*e*. time to commence mating (duration from the mounting to intromission of aedeagus), and mating duration (time from intromission unt il dismounting), were observed. After mating females were placed in separate Petri dishes containing *ad libitum* supply of aphid for egg laying. Before the oviposition period, fecundity (the number of eggs laid) and egg viability were recorded for next seven days.

## Statistical analysis

Data on time to commence mating, mating duration, fecundity, pre-oviposition period and egg viability (independent factors) was first tested for normality and homogeneity using Shapiro-Wilk’s method and Levene’s method, respectively. Data were found to be non-normal (time to commence mating: F = 0.715, df = 90, P < 0.05; mating duration: F = 0.875, df = 90, P < 0.05; pre-oviposition period: F = 0.766, df = 90, P < 0.05; average fecundity: F = 0.802, df = 90, P < 0.05; average percent egg viability: F = 0.751, df = 90, P < 0.05) and non-homogenous (time to commence mating: F = 56.751, df = 2,87, P < 0.05; pre-oviposition period: F = 33.479, df = 2,87, P < 0.05; average fecundity: F = 27.945, df = 2,87, P < 0.05; average percent egg viability: F = 22.681, df = 2,87, P < 0.05) except for mating duration (F = 1.498, df = 2,87, P > 0.05). Mating did not occur in the first heterospecific mating combination between *P. dissecta* male with *C. sexmaculata* female (Treatment b). Therefore, data of this treatment were excluded from analysis.

Data on mating (Time to commence mating and mating duration) and reproductive parameters (pre-oviposition period, fecundity and egg viability) were subjected to univariate analysis using a GLM. The data on reproductive parameters (fecundity and percent egg viability) were subjected to an Analysis of covariance (ANCOVA) with mating duration as a covariate. A Bonferroni post hoc test was used to determine the level of significance in case of significant main effects. All the analyses were performed on SPSS Version 20 and all plots were made using SPSS Version 20.

## Results

### Mating parameters

Time to commence mating (TCM) in *P. dissecta* was significantly affected by the presence of heterospecific adults (F = 5.733, P < 0.05, df = 2,89). The longest TCM was recorded in heterospecific treatment while shorter TCM’s were recorded in control (male and female of *P. dissecta*) and mixed treatment (both male and female of *P. dissecta* with *C. sexmaculata* male), respectively (Figure 1).

**Figure 1:**
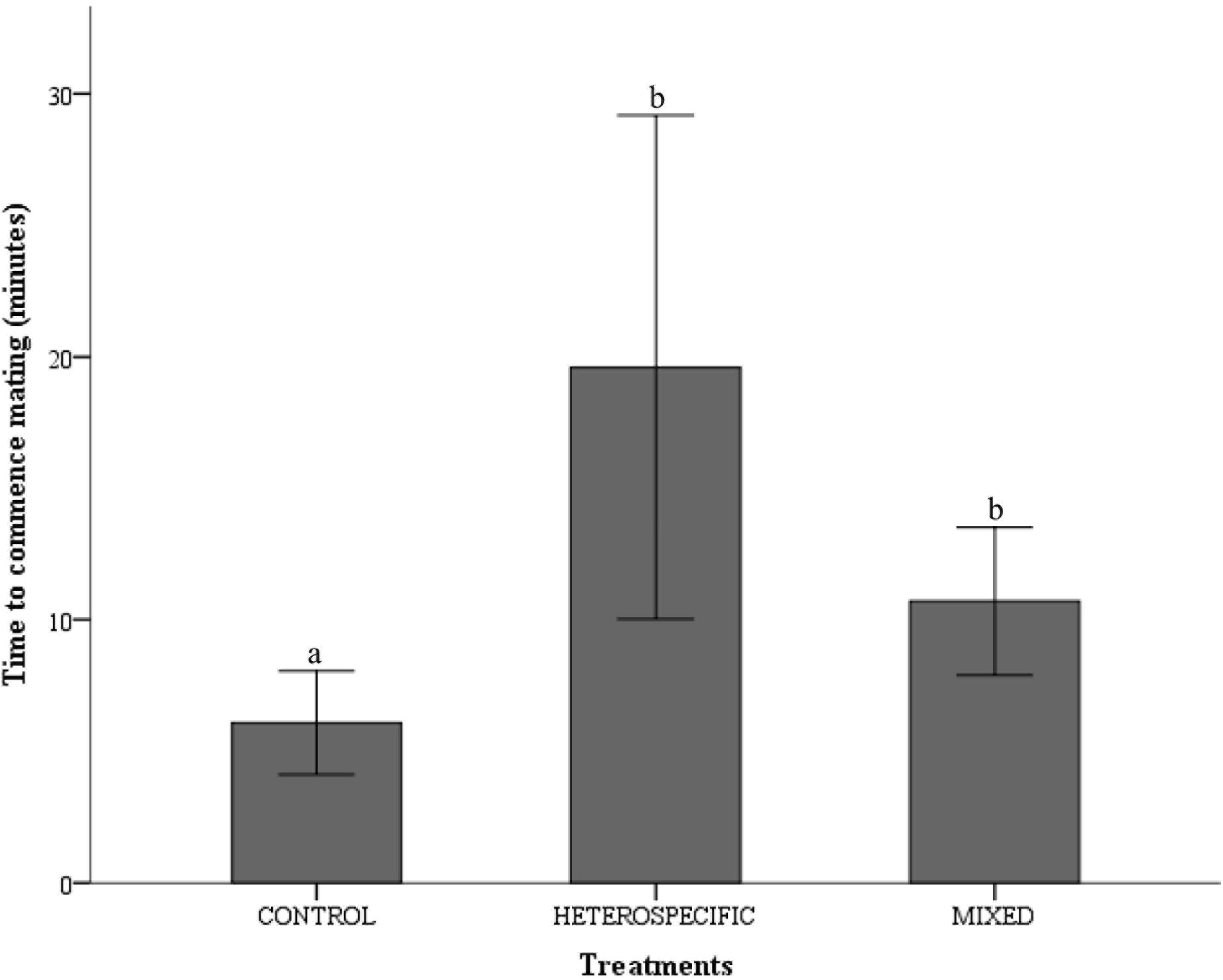
Time to commence mating (TCM) in different mating treatments *i*.*e*. control (*P. dissecta* male × *P. dissecta* female), heterospecific (*C. sexmaculata* male × *P. dissecta* female), and mixed (*P. dissecta* pair × *C. sexmaculata* male) mating treatments.

Mating duration (MD) was found to be significantly affected by the presence of heterospecific adults (F = 138.077, P < 0.05 and df = 2,89). The longest MD was recorded in control and mixed treatment while the shortest MD was recorded in heterospecific mating treatment (Figure 2).

**Figure 2:**
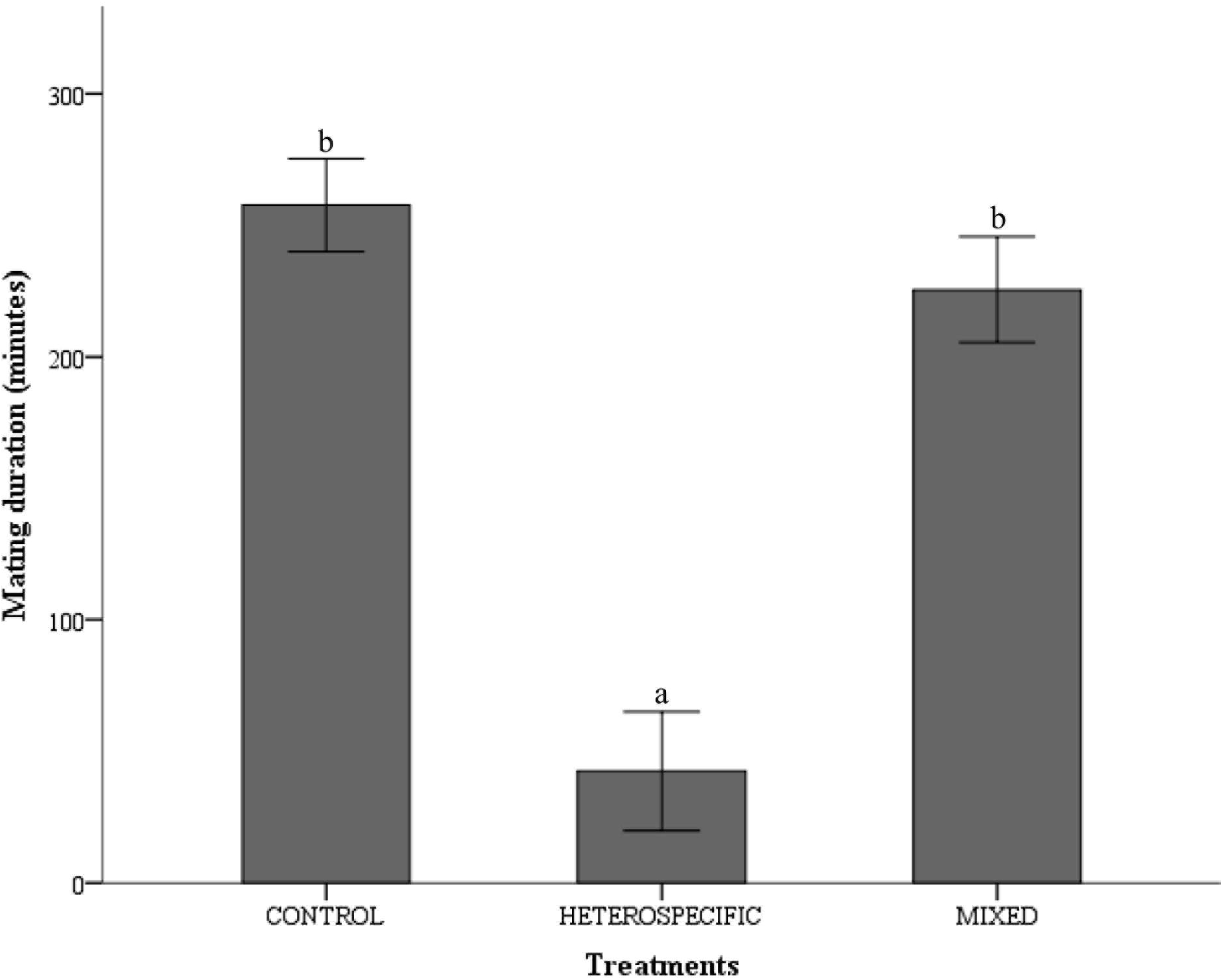
Mating duration (MD) in *P. dissecta* in different mating treatments *i*.*e*. control (*P. dissecta* male× *P. dissecta* female), heterospecific (*C. sexmaculata* male × *P. dissecta* female), and mixed (*P. dissecta* pair × *C. sexmaculata* male) mating treatments.

We observed conspecific mating in all replicates (30 out of 30) of the control and mixed treatments, while in the heterospecific treatment, heterospecific mating did not occur in the first heterospecific mating combination with *P. dissecta* male and *C. sexmaculata* female (0 out of 30); however, in second combination of heterospecific mating, interspecific mating was observed in *C. sexmaculata* male and *P. dissecta* female in 43.33% replicates (13 out of 30).

### Reproductive parameters

The pre-oviposition period in *P. dissecta* was significantly affected by the presence of heterospecific adult (F = 18.355, P < 0.05 and df = 2,89). The shortest pre-oviposition period was recorded in control treatment and the longest pre-oviposition period was recorded in mixed treatments, but they were not statistically significant. A significant difference was observed due to heterospecific treatment since *P. dissecta* females of the heterospecific mating treatment did not lay any eggs (Figure 3).

**Figure 3:**
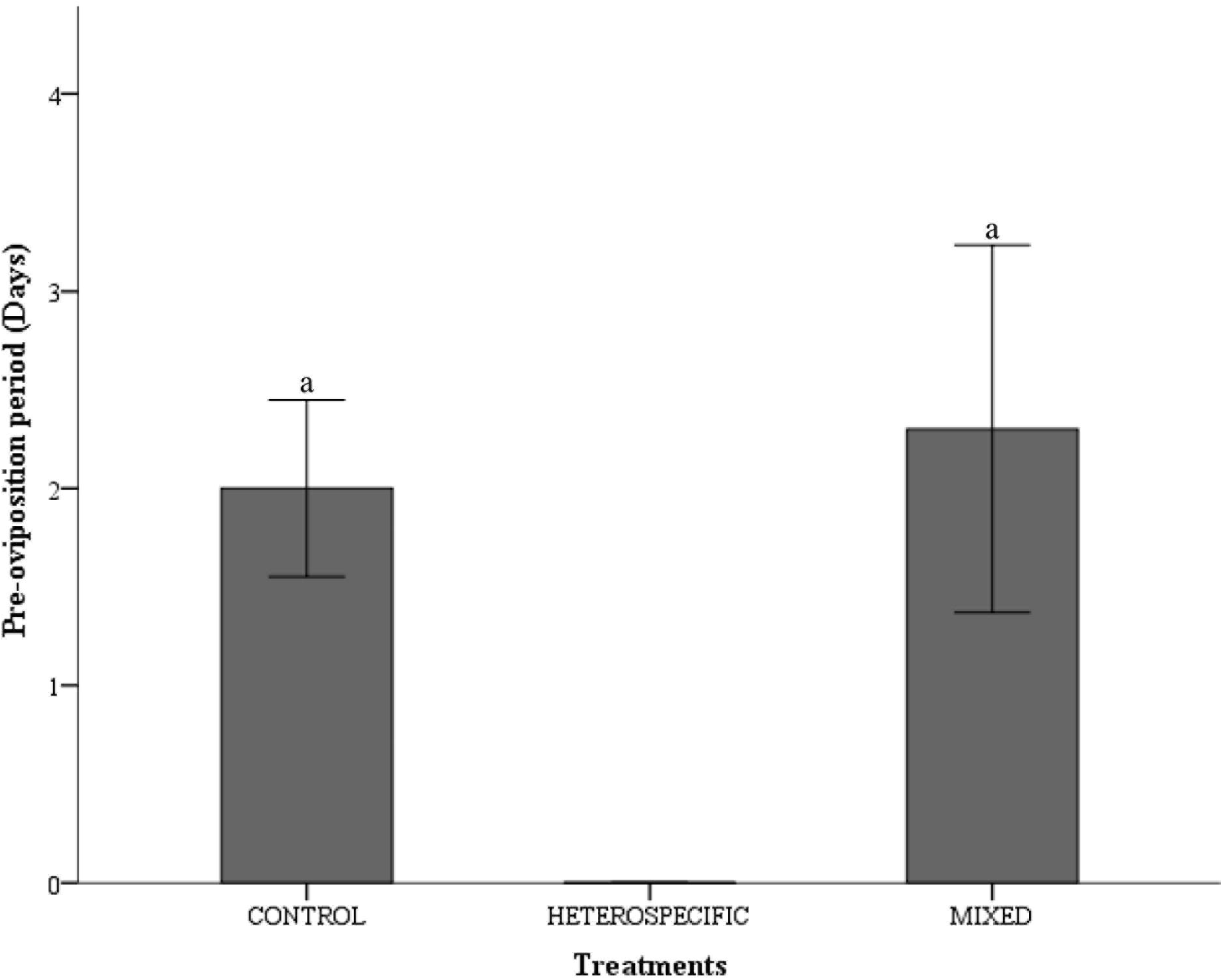
Pre-oviposition period of different mating treatments *i*.*e*. control (*P. dissecta* male× *P. dissecta* female) and mixed (*P. dissecta* pair × *C. sexmaculata* male) mating treatments.

Fecundity in *P. dissecta* was significantly affected by the presence of heterospecific adult (F = 35.449, P < 0.05 and df = 2,89). Females of the control group were more fecund than those of the mixed treatment (Figure 4).

**Figure 4:**
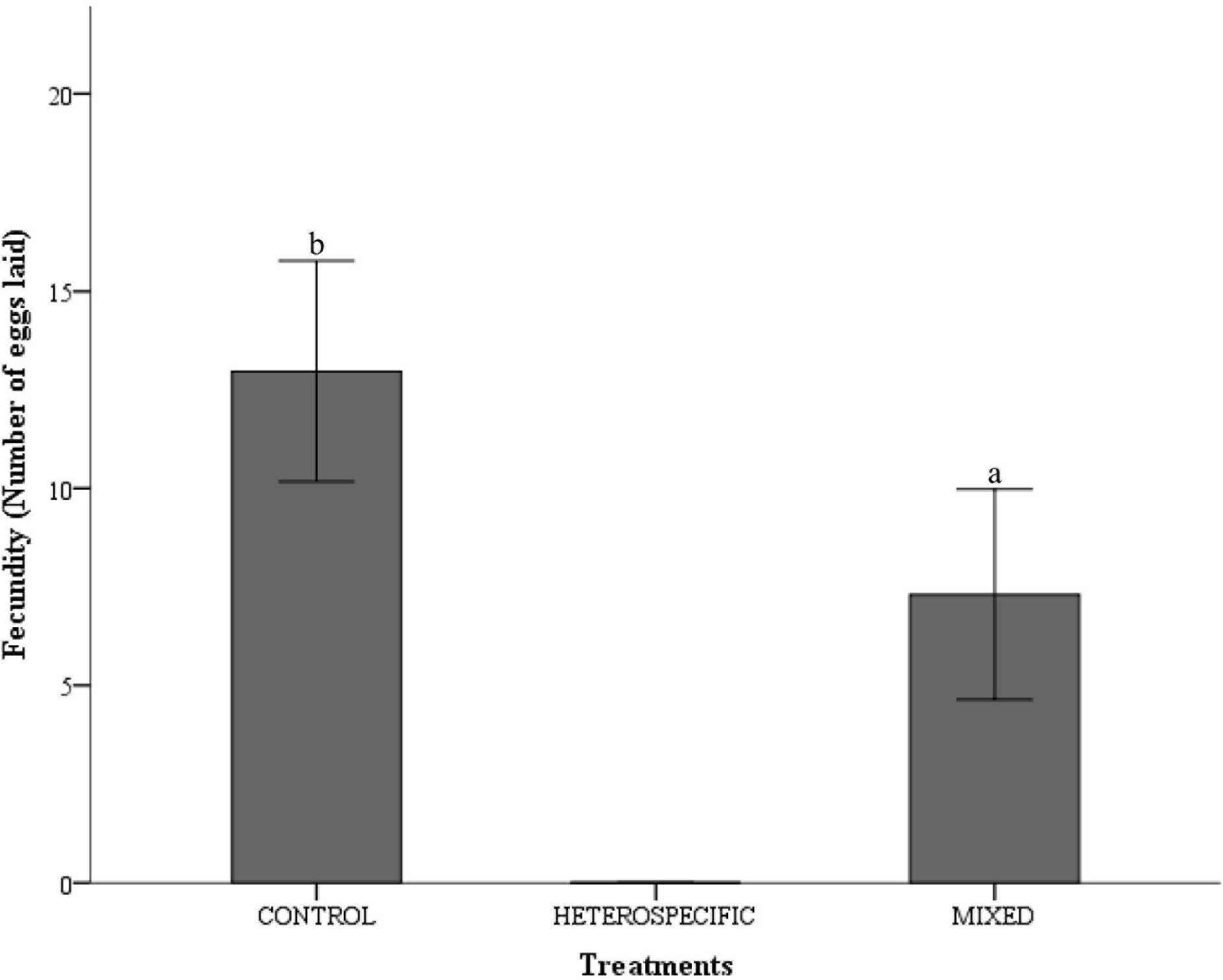
Average fecundity of females in control (*P. dissecta* male× *P. dissecta* female) and mixed (*P. dissecta* pair × *C. sexmaculata* male) mating treatments.

Egg viability in *P. dissecta* was significantly affected by the presence of heterospecific adults (F = 79.224, P < 0.05 and df = 2,89). The lowest percentage of egg viability was recorded in mixed treatment compared to control treatment (Figure 5). However, when fecundity and egg viability were subjected to an ANCOVA with mating duration as a covariate, an insignificant effect of heterospecific interference on fecundity (F = 1.835, P > 0.05, df = 2,89); egg viability (F = 0.881, P > 0.05 and df = 2,89) and mating duration (Fecundity: F = 0.892, P > 0.05 and df = 1,89; percent egg viability: F = 2.182, P > 0.05 and df = 1,89) was observed.

**Figure 5:**
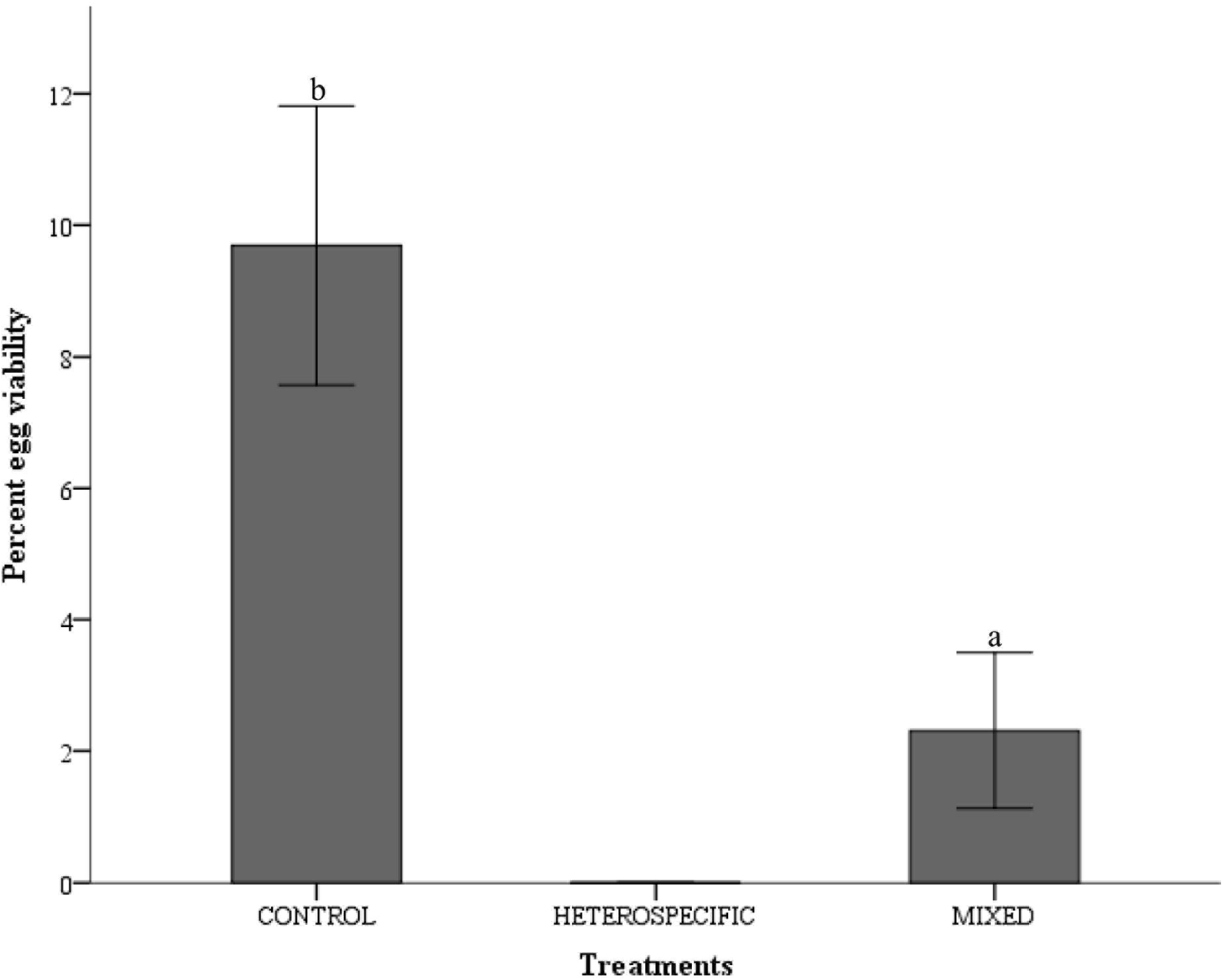
Percent egg viability in control (*P. dissecta* male× *P. dissecta* female) and mixed (*P. dissecta* pair × *C. sexmaculata* male) mating treatments.

## Discussion

Our results from laboratory experiments revealed that heterospecific mating occurred between *P. dissecta* females and *C. sexmaculata* males; however, no mating was recorded between *C. sexmaculata* females and *P. dissecta* males. Mating parameters (time to commence mating and mating duration) were found to be significantly affected by the presence of heterospecific mates. Shorter times to commence mating were recorded in control and mixed treatments while the longest time to commence mating was recorded in heterospecific treatments. In addition, longer mating durations were recorded in control and mixed treatments, and the shortest mating duration was recorded in heterospecific treatments. The reproductive parameters (pre-oviposition period, fecundity and percent egg viability) were also significantly affected with highest fecundity and percent egg viability in control treatment and lowest in mixed treatment.

Longer durations to commence mating were recorded in heterospecific (male *C. sexmaculata* and *P. dissecta* female) and mixed (with both males and females of *P. dissecta* and *C. sexmaculata* male) mating treatments. This may be attributed to the interference caused by heterospecific species in both heterospecific and mixed mating treatments. Shorter time to commence mating in control treatments consisting of conspecifics mates may be attributed to their mate recognition ability and genital compatibility. In *Allonemobius* species, longer time to commence mating (first mount) has been reported in heterospecific mating (Birge *et al*., 2010).

In the first heterospecific combination, multiple contacts between *P. dissecta* male and *C. sexmaculata* female occurred but mating did not commence. While in the case of *C. sexmaculata* male and *P. dissecta* female, the male mounted over the female but time to commence mating increased which might be attributed to the fact that the mate possibly took longer to recognise the conspecifics versus heterospecifics and failed in species recognition possibly owing to their close phylogenetic relationship and similarity of volatile hydrocarbons (Pattanayak *et al*., 2014; Pattanayak *et al*., 2016) In some cases, *P. dissecta* females also showed rejection behaviour by running around in the Petri dish and *P. dissecta* males showed leg movement during interference. This type of behaviour is usually related to defensive behaviour in family Tetrigidae (Orthoptera) and includes leg movements and body shaking (Hochkirch *et al*., 2007).

Furthermore, the shortest mating duration was recorded in heterospecific treatment due to different species. In mixed treatment; heterospecific adult male (*C. sexmaculata*) disturbed *P. dissecta* male and female during mating and showed reproductive interference by making mate attempts with *P. dissecta* female and climbed either on male or female of *P. dissecta*. Reproductive interference is responsible for decreasing mating duration in mixed treatment.

In *Aedes* mosquito species, mating duration has been reported to be significantly affected by heterospecific mating (Zhou *et al*., 2022). In *Tetrix* species (Ground-hoppers), a longer mating duration was also recorded in the control treatment and the shortest mating duration was recorded in mixed treatment (Hochkirch *et al*., 2007).

A similar study in *Drosophila* species also reported a shorter mating duration in the case of conspecific mating (*D. persimilis* female with *D. persimilis* male) and a longer mating duration in heterospecific mating (*D. persimilis* female with *D. subobscura* male) (Manzano-Winkler *et al*., 2017).

The pre-oviposition period was also found to be significantly affected with a similar pre-oviposition period in control and mixed treatments; however, no egg laying was observed in heterospecific mating combinations which might be attributed to postmating barriers or the absence of fertilization. In mosquito, *Aedes* egg laying was not observed in heterospecific mating (Paton & Bonsall, 2019; Zhou *et al*., 2022). Thus, ultimately heterospecific mating results in loss of gametes, energy and time (Gröning & Hochkirch, 2008; Burdfield-Steel & Shuker, 2011).

The lowest fecundity and egg viability were recorded in mixed treatments due to presence of heterospecific mates and reproductive interference; while in a heterospecific treatment the females did not lay eggs. After mating, in a mixed treatment, we observed that heterospecific adult (*C. sexmaculata* male) copulate with *P. dissecta* female many times and interrupted egg laying. Similar results were also reported in Ground-hoppers (*Tetrix*), Seed beetles (*Callosobruchus*) (Kyogoku & Nishida, 2012; Kishi & Nakazawa, 2013; Kishi, 2015), and Squash bugs (*Anasa*) (Villa *et al*., 2021). Studies in *Aedes* mosquitos have also reported that the females do not lay eggs after heterospecific mating (Zhou *et al*., 2022; Paton & Bonsall, 2019). In contrast, some insects are known to produce viable and hybrid offspring after heterospecific mating, such as whiteflies (*Bemesia* sp.) (Crowder *et al*., 2010), and ladybirds (*Coccinella* sp.) (Noriyuki & Osawa, 2016). In some cases, inviable and hybrid eggs have been reported, such as copepods (*Skistodiaptomus* sp.) (Thum & Ryan, 2007). A low egg viability was also recorded in bumble bees (*Bombus* sp.) (Tsuchida *et al*., 2019).

Reproductive interference has been commonly observed under laboratory conditions, but it is not always found in field conditions (Orr, 1989; Thum & Ryan, 2007). In field conditions, *P. dissecta* and *C. sexmaculata* coexist together and they have same mating signals, body size and same life history. Therefore, chances of heterospecific mating are high but eggs are not produced due to the species belonging to different genera. This asymmetric mating-induced infertility in the female is considered responsible for species displacement (Zhou *et al*., 2022).

Our results support the hypothesis that reproductive success of *P. dissecta* was significantly affected by the presence of heterospecific adults. Our results show that *C. sexmaculata* males are more active and polygynous insects compared to *P. dissecta* males and *C. sexmaculata* males attempted heterospecific mating more frequently.

This study concludes that: (1) heterospecific mating did not commence between *P. dissecta* male and *C. sexmaculata* female, while heterospecific mating was recorded between *P. dissecta* female and *C. sexmaculata* male, (2) *C. sexmaculata* males interfere with the conspecific mating of *P. dissecta* through multiple mating attempts and reduced their fecundity and percent egg viability.

## Acknowledgements

MS gratefully acknowledges CSIR, New Delhi, India, for Junior Research Fellowship, (09/107(0416)/2021-EMR-I) dated April 8, 2021.

MS is also thankful to Mr. Nick Bailey, Department of Biological Sciences, Auburn University, Alabama, United States, for improving the language of the manuscript.

